# The 5’ regulatory region of the β actin gene in *Clarias* species is complex and variable in relation to ecological needs

**DOI:** 10.1101/2020.05.13.093930

**Authors:** Deepali Sangale, Anita Tiknaik, Gulab Khedkar, Danid Haymer, Chandraprakash Khedkar, Shrish Tiwari

## Abstract

The β actin gene is involved in various cellular housekeeping processes including transcription, mRNA processing, cell signaling and chromosome remodeling. For regulating the expression of this gene under different environmental conditions, the promoter region of the β actin gene is structurally dynamic with multiple regulatory features in the upstream region. Most previous information about the 5’ regulatory region of the β actin gene has been limited to *in vitro* laboratory experiments. Considering the need for functional versatility of expression of this gene in the Catfish *Clarias batrachus* in different environments, here we have analyzed the 5’ regulatory region of β actin and identified numerous elements that are variable. We have made comparisons of individuals from three populations found in three different diverse ecological systems, as well as in three sister species, to elucidate its structural diversity. Our results show that the 5’ regulatory region has considerable diversity and changes in architecture with respect *Cis*-acting regulatory elements. These changes may be linked to positive selection in combating pollution or disease like conditions encountered by the organism. These observations leads to the conclusion that 5’ regulatory region of a housekeeping gene like β actin, modify its architecture as per the environmental conditions. These modifications specifically includes diversity of TF binding sites indicating the assortment of environmental variables and only one third region of 5’ regulatory region is conserved which was yet not highlighted.

**Author summary:** Promoter is a regulatory region where the basal transcription machinery assembles to initiate the process of transcription. It plays crucial role in controlling the gene expression. The 5’ regulatory region includes TATA box, CAAT box, GC box and Cis -acting regulatory elements. Most previous information about the 5’ regulatory region of the β actin gene has been limited to in vitro laboratory experiments. Our study results show that the 5’ regulatory region has considerable diversity and changes in architecture with respect *Cis*-acting regulatory elements. These changes may be linked to positive selection in combating pollution or disease like conditions encountered by the organism. These observations leads to the conclusion that 5’ regulatory region of a housekeeping gene like β actin, modify its architecture as per the environmental requirements. These modifications precisely includes diversity of TF binding sites indicating the assortment of environmental variables and only one third region of 5’ regulatory region is conserved. These findings clearly define a novel role of promotor of β actin gene which was yet not highlighted. These findings can broaden our understanding in linking TF in 5’ regulatory regions to a specific environmental variable/disease conditions. This may become a simple strategy in understanding complex gene-environment interactions.

## Introduction

Actin proteins are among the most vital and abundant cytoskeletal elements of a cell. They play critical roles in a wide array of cellular processes including cell division and the regulation of gene expression. These functions are ascribed to the ability of actin to form filaments that can promptly assemble and disassemble rendering to the needs of the cell. In vertebrates, there exist six different but exceptionally conserved actin isoforms [1–4]. Each is the product of a separate gene, with *Actb* and *Actg1* encoding for the abundantly expressed β-actin and γ-actin cytoplasmic isoforms, respectively.

The coding regions of these genes are highly conserved in diverse organisms. In birds and mammals, for example, β-actin and γ-actin differ at only four biochemically similar amino acid residues. This has been interpreted as signifying evolutionary pressure through environmental selection to maintain these apparently minor sequence differences. Recently it has also been established that these amino acid differences confer unique biochemical properties between the two isoforms [5, 6]. Understanding how these distinct properties relate to functional differences within the cell, however, remains elusive.

β actin is a major cytoplasmic actin isoform that is conserved and constitutively expressed in non-muscle eukaryotic cells as a housekeeping protein [7]. It plays an important role in maintenance of the cell cytoskeleton, cell migration, cell division, cell junction formation and vesicle trafficking [8, 9].β actin is also involved in key nuclear processes such as transcription, mRNA processing, cell signaling and chromosome remodeling [10–12]. In addition, studies have shown that the regulatory region of the β-actin gene is highly efficient in regulating the expression of downstream foreign genes [13]. For these and other reasons, β-actin is widely used as internal marker or reference gene for studies quantifying gene expression [14]. However, information about the 5’ regulatory region of the β actin gene is still limited to *in vitro* laboratory experiments which do not adequately take into consideration gene-environment interactions. For variation in natural populations, however, it is crucial to comprehend this phenomenon.

To better understand this functional versatility, here we have analyzed the 5’ regulatory region of the β actin gene of the catfish *Clarias batrachus.* We identified a 5’ *Cis*-acting element and CpG islands which are essential for regulatory activity of this region. The binding activity of the 5’ *cis*-acting elements can be found in a wide variety of tissues, and this may play a role in the transcriptional activation of numerous promoter elements containing specific factor recognition sites. CpG islands, for example, have been predicted to play dual roles in 5’ regulatory regions and local chromatin structuring [15]. As part of our study, we compared the 5’ regulatory region of this gene from three populations of the same fish found in three different diverse ecological systems (a dam, the rivers and a hatchery) to correlate its variability in response to ecological pressures. We also made comparisons to the same gene in three sister species (*C. gariepinus, C. magur* and *C. dussumieri*) to further document structural diversity.

## Results and Discussion

To our knowledge, this study is the first report on characterization of a β actin promoter from *C. batrachus,* a commercially important species used in aquaculture. The sequence data were deposited in NCBI GenBank under BioProject SRA accession No. **PRJNA623463** which can be accessed through the link https://www.ncbi.nlm.nih.gov/sra/PRJNA623463.

### Organization of the β actin promoter

The 5’ regulatory region of this gene, consisting of 1262 bp, was successfully amplified for characterization. This is a simple and robust method for producing multiple copies of target sequences [16] compared to other cloning methods [17]. The amplicon derived from this gene was sequenced and characterized, including annotation of the complete assembly of 5’ regulatory region (Fig 1.) The sequence of this region of *C. batrachus* was also compared with 5’ regulatory regions from a sister species, *C. gariepinus* (GenBank accession no. **EU527190.2**). Functional annotation by comparison between closely related or sibling species is often used as a first step to identify conserved regions vs. sequences that have changed during evolution [18].

**Fig 1.**
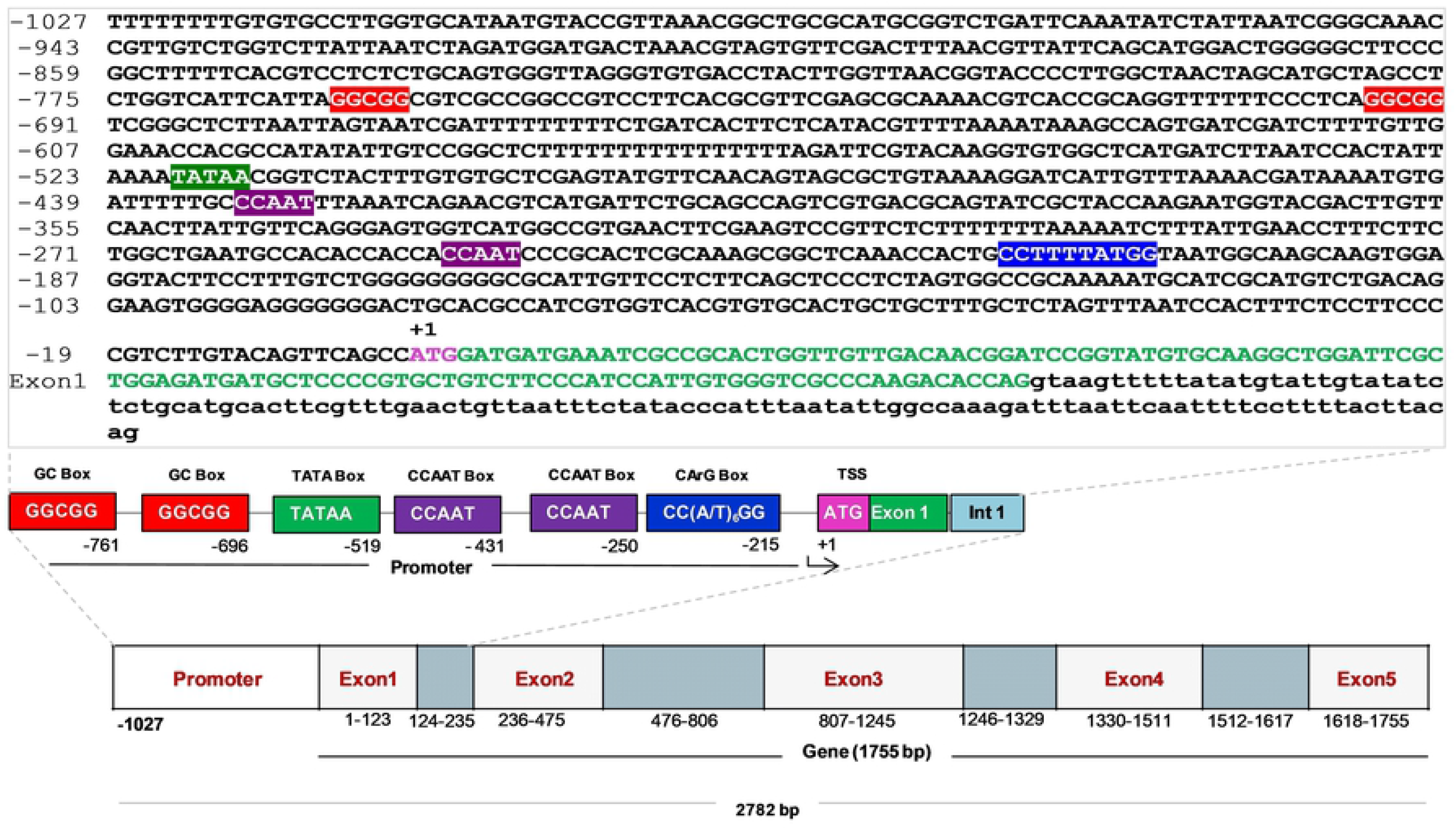
Schematic map showing the genomic organization of β actin gene and its 5’ control region from habitat “B” (2782 bp) (Predicted transcription start site (TSS) is denoted with +1. First base of starting codon ATG was set as positon +1. The Exons (upper case letters in green color) and introns (lower case letters in black color) are shown in different boxes. *Cis*-elements including TATA Box, CAAT Box, CArG Box and GC Box are denoted in the figure panel. The number below the Exon, Intron and *Cis*-element boxes indicate the position of nucleotides. Numbering for the nucleotide sequence position is given on left side, underlined letters shows the TATA, CAAT, CArG and GC boxes and the numbers above it shows the positions of regulatory elements)

In our study the 5’ regulatory region of this gene in *C. batrachus* from habitat “A” consisted of a total of 1262 bp. This extends from base position −1027 to −1 bp of the promoter region upstream of the ATG initiation codon oftranslatedexon1 (123 bp) and intron 1 (111 bp) (Fig 1). The regulatory elements in this region included a conserved TATA box (TATAA) predicted at position −519, two CAAT boxes (CCAAT) predicted at positions −250 and −431. A consensus CC(A/T)_6_GG motif (CArG box: CCTTTTATGG) was identified at position −215 along with two GC box at positions −696 and −761 upstream from the initiation codon ATG at +1 site (Fig 1). Of the two CAAT boxes present, only one may be an important regulatory factor leading to high levels of β actin gene transcription [8]. Other studies reporting the existence of more than one CAAT box suggests that they may be involved in the up-regulation of a number of diverse promoters [19, 8].

### Similarity search for assembled β actin promoter

Sequence similarity searches were conducted to validate the β actin promoter assembly built here. These results indicated relatively high levels of sequence similarity with other species, but no region of this gene showed more than 85% similarity. Outside of this region, the level of query coverage was very low (<10%) (S2A Table). The result showing that sequences upstream of the 5’regulatory region of the β actin gene are less conserved has also been observed for other housekeeping type genes [20–22].

### Features of Cis-acting regulatory elements (CAREs)

Predictions were made for the locations of the binding sites for *Cis*-acting regulatory elements. These included motifs for the specificity protein1 (Sp1 Krupel like factor), the cAMP Response Element (CRE), Nuclear Factor-1 (NF-1), a myogenic regulatory factor (Myf), Activator Protein-1 (AP-1), an E-box family motif of the proto-oncogene (Myc), a late SV40 Factor (LSF), an E26 transformation specific family (Ets) sequence, and a Serum Response Factor (SRF), also known as a CArG motif (Fig 2). *Cis*-acting regulatory elements such as these, are known to provide sites for DNA binding proteins to modulate the expression of gene at both transcriptional and post transcriptional levels [23]. Because of this, the prediction and functional characterization of *Cis*-acting elements is a crucial step in understanding the precise mechanisms of gene regulation [24, 25].

**Fig 2.**
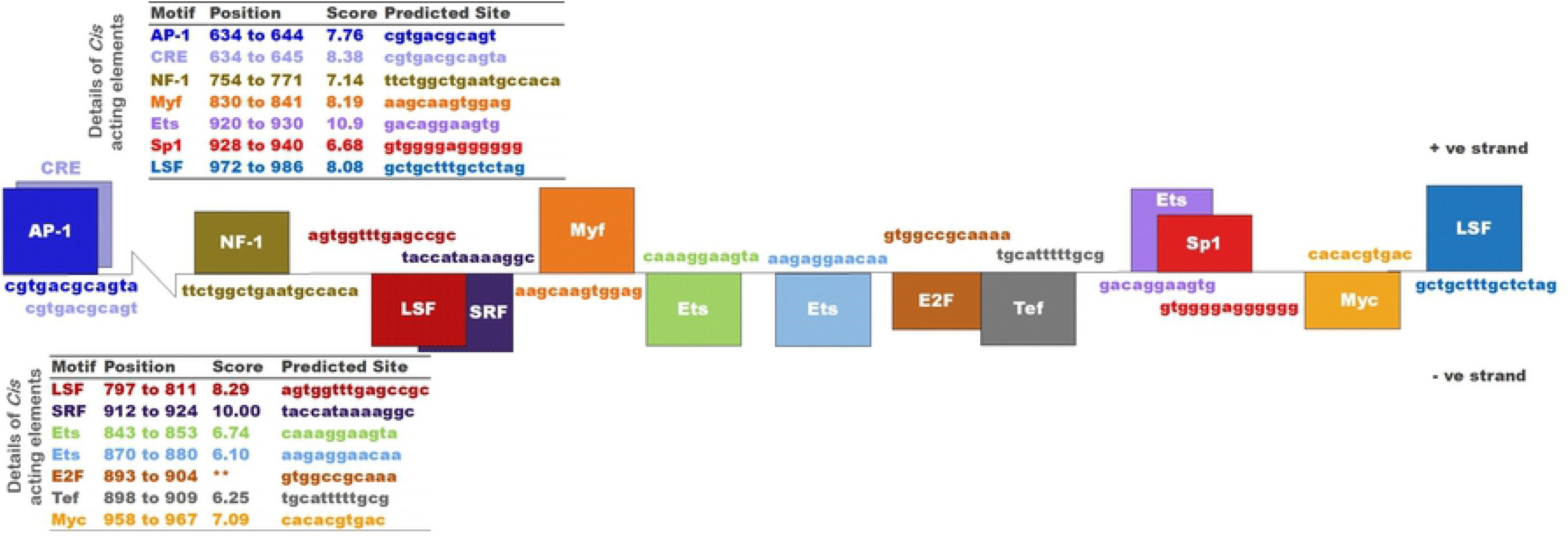
Diagrammatic representation of *Cis*-acting element and their binding sites in β actin promoter. (The boxes on upper and lower side indicate the Cis-element and the sequence show the binding site for each element on +ve and –ve strand respectively. The table indicates the position and score of each element. ** -no score is predicted with Cister software).

The ultimate goal of any study of these elements is to ascertain the specific role of each factor in control of transcription regulation and fundamental processes such as cell growth, division, differentiation, migration, survival during stages such as embryonic development and transcription regulation (Table 1) [51–53, 3, 10].

**Table 1.**
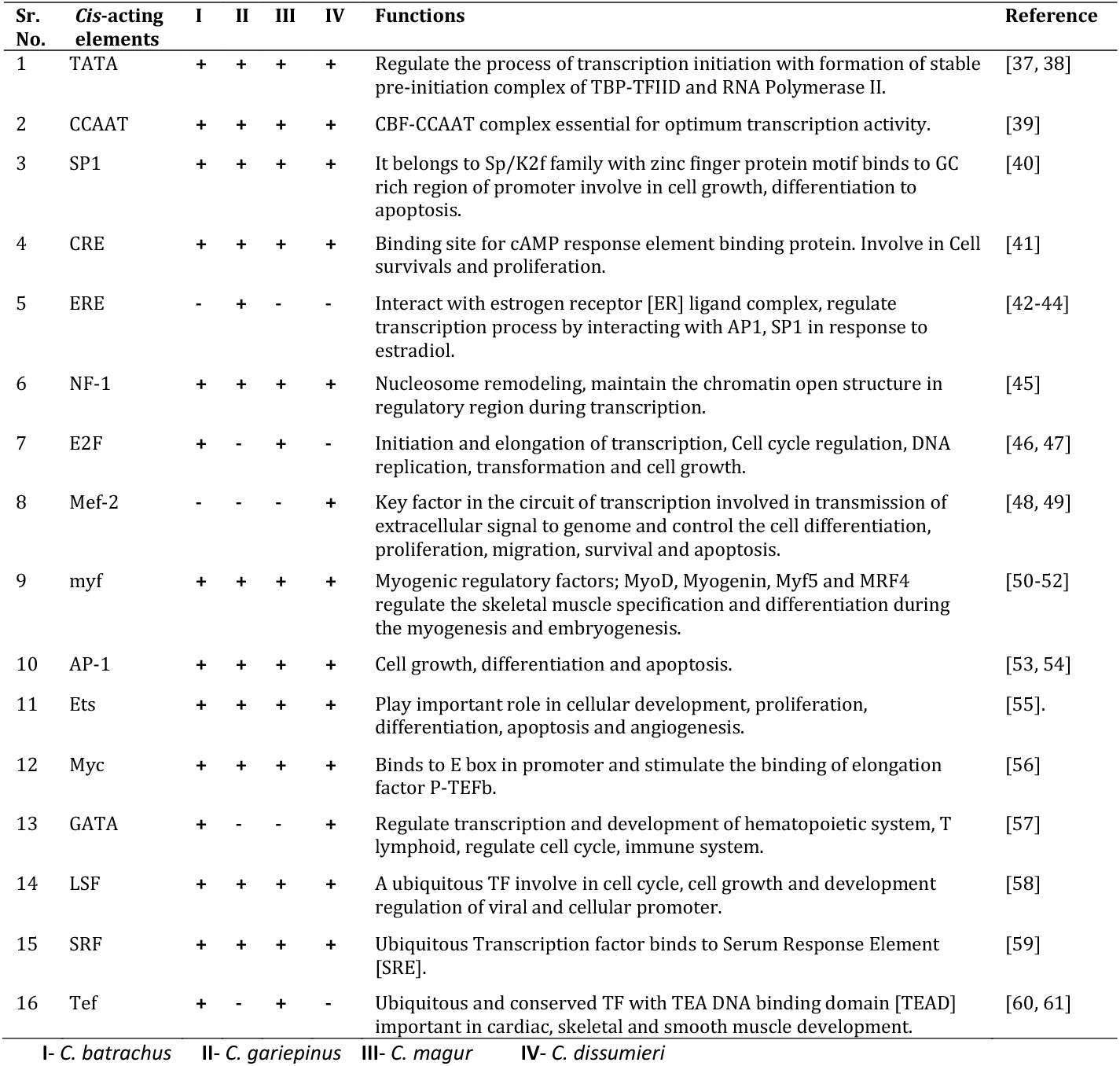
Comparison of *C/s*-acting elements among four species of genus Clarias.

In addition, though, the β actin gene of a species like *C. batrachus* can also be used as an internal control gene or reference housekeeping gene (54-57).

However, other factors like Mef-2 and Ets also appear to be involved in processes like apoptosis and angiogenesis, respectively. Ruan and Lai [58] and Guo *et al.,* [59] also noted that expression of the β actin gene is unstable in response to biochemical stimuli during growth and differentiation in disease conditions like cancer. These insights about the β actin gene may ultimately overshadow its role as an internal marker or reference gene in gene expression studies [60–64].

We also compared the transcription factor binding sites in the 5’ regulatory region of this gene in *C. batrachus* to those of sister species such as C*. gariepinus, C. magur* and *C. dussumieri* (Table 1). From the comparisons made, it was evident that except for five *Cis*-acting elements (ERE, E2F, Mef-2, GATA and Tef), all others were common in species within the genus Clarias. The ERE and Mef-2 were absent in all three populations of *C. batrachus* studied here. This suggests that the Clarias β actin gene characterized here can be treated as reference gene for expression studies in *C. batrachus*, but not for other species in the genus studied here.

### Comparison of 5’ regulatory region among the species belong to different habitats

The expression of any gene may be extensively influenced by factors present in local environments [65–67] including conditions such disease, pollution, competition, etc. [68–70]. In an effort to delineate that structural diversity may play in environmental response [22]. We characterized the regulatory region of this gene in multiple individuals of *C. batrachus* from three different habitat as shown in supplementary table S1.

The multiple sequence alignment of the 5’ regulatory regions of β actin gene for individuals of *C. batrachus* from three habitats *viz.* habitat “A” (Dam population), habitat “B” (River population) and habitat “C” (Hatchery population) show differences in the number of nucleotides (Habitat A=1027 bp, Habitat B=1586 bp and Habitat C=1572 bp) (Fig 1, S2 and S3 Figs). Other specific differences include the absences of the untranslated exon 1 (100 bp), one TATA box, two CCAAT boxes and one CArG motif (SRF) in the 5’ regulatory region of the β actin gene of the fish from habitat “A” (Table 2 and Fig 3).

**Fig 3.**
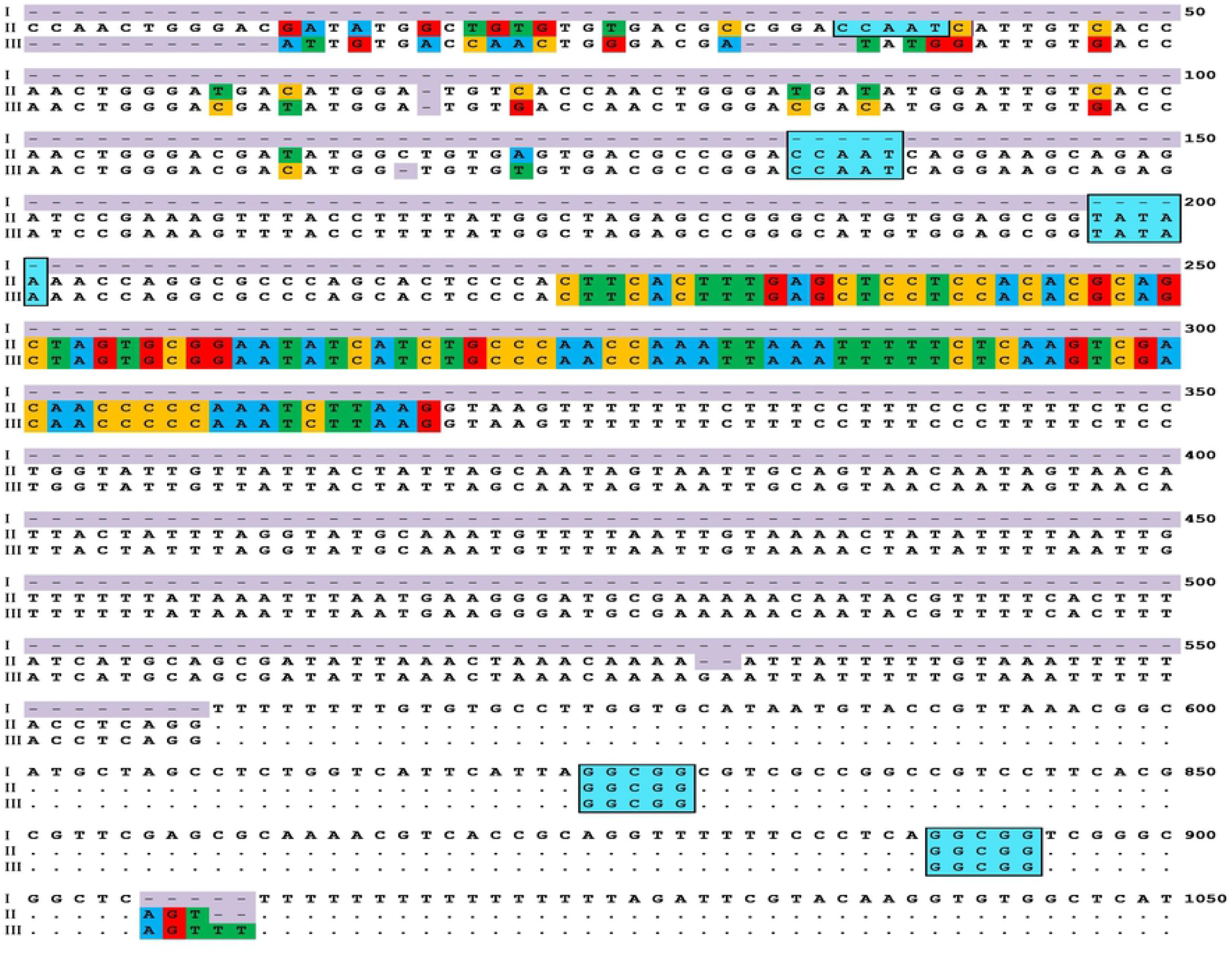
Sequence alignment and comparison of β actin promoter of *C. batrachus* from different habitats. I. Nucleotide sequence of *C. batrachus* collected from a dam site (Habitat “A”). II. Nucleotide sequence of *C. batrachus* collected from a river site (Habitat “B”). III. Nucleotide sequence of *C. batrachus* collected from a hatchery (Habitat “C”). (“.” indicates the identical nucleotides at respective position; “-” are gaps introduced for optimal alignment; the close boxes marks the conserved sequences and factor binding sites, CAAT box, TATA box and GC box respectively. The nucleotide sequence shading in color indicate the variable sites and untraslated exon 1 is present in 250-350bp fragment of the promoter region. The numbers on right side indicate the nucleotide position.

**Table 2.**
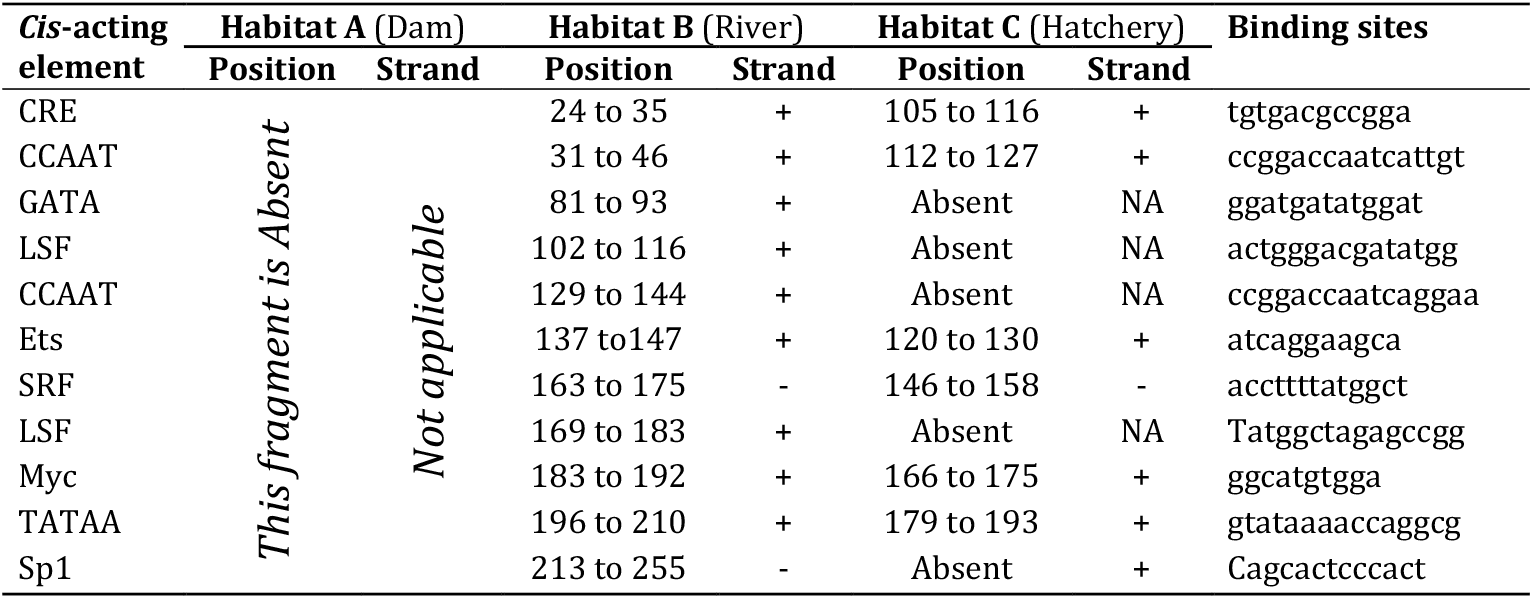
Mapping *Cis*-acting element among different habitats.

However, the structural organization of the various vertebrate genes (for e.g. GATA gene) shows numerous common features, including the use of multiple variably distant enhancers and the presence of alternative 5’ untranslated exons whose expression is regulated by distinct, and often tissue-specific, promoter units [71]. Also, 25 variables sites were seen in the first 150 bps of the 5’ regulatory region of the β actin gene in the fish from habitat “B” and habitat “C”. This accounts for 16.6% of the variation (Fig 3). Additionally, five variable sites were observed in the region starting from 1006-1010 bp, surprisingly, this region was absent in the fish from habitat “A” whereas, two sites (1009-1010) were absent in fish from habitat “B” (Fig 3).

However, the sequences in the remaining region of 582 bp (1011 bp to 1593 bp) are completely conserved in the fish from all three habitats studied here. This is relevant to the observations by Tarallo et al., [72] where it was noted that environmental factors affect not only on physiology and morphology but can also exert an influence on the entire genome composition due to selection pressure. In such cases, the effect of environmental factors is mainly seen on the GC content of specimens within each habitat [73–75].

In our study, we selected three individuals from different habitats for characterization of 5’ regulatory regions to consider in light of the finding made by Foerstner et al., [75] and Tarallo et al., [72] (S3 Table). From our results, we can conclude that natural selection and adaptive evolution is having an impact on the nucleotide diversity in the regulatory sequences or TF binding sites. Further, it can be concluded that such changes in nucleotide diversity are associated with positive selection leading to resistance to external stimuli such as pathogens or environmental pollutants [76]. However, some population level variations in transcription factors were noticed (Table 2), as indicated in habitat “A” where binding sites for all eleven *Cis* acting elements (CRE, CCAAT (2 nos.), GATA, LSF, Ets, SRF, LSF, Myc TATAA and Sp1) were absent. This suggests that for the individuals in the population at habitat “A”, their promoter architecture has evolved through deletions of binding site recognition sequences. This may also be part of an effort to adapt rapidly to environmental selection pressures as suggested by Maury and Marguerat [77]. In such situations, the strength of the transcription factor binding, as reflected in the number and positioning of binding sites may strongly affect the level of gene expression in response to changed environmental conditions [78, 79].

Data from Table 2 and Supplementary Fig S2 and Supplementary Fig S3 also clearly indicates that individuals in the population from habitats “B” and “C” have more TF binding sites, and the complex nature of this promoter region may be related to adapting to drastic environmental conditions [80]. Several authors noticed that the River Ganga (Habitat B) is one of the highly polluted river systems in India [81, 82], and this might impose extreme selection pressure on the fish. As evidenced in this study, in this habitat the promoter region has undergone functional changes which may be related to this pressure [83]. In comparison, the β actin promoter architecture of fish from habitat “C” was found to be less complex (S3 Fig). This may be correlated with the fact that when rivers in India were ranked for several pollution parameters, the River Ganga ranked at this first position in terms of pollution levels whereas the River Godawari, the site for habitat “C” is indicated to be only moderately polluted [84–87].

Overall, these results suggest that environmental factors impact the evolution of 5’ regulatory region with respect to presence of specific regions such as binding sites for TFs in the β actin gene. In our analysis, the distribution of number of TFs per promoter varies in different environments. Promoters with high densities of TF suggests that they may require complex regulation to be able to adapt to specific but variable environment factors (such as pollution).

### Comparison of Untranslated exon (*in silico*)

Comparison of the untranslated exon of *C. batrachus* with other teleost and mammalian species such as human and mouse is shown in Fig 4. The alignments indicate that the 5’ regulatory region of *C. batrachus* β actin gene from habitat “A” does not contain an untranslated exon. However, in the *C. batrachus* populations from habitats ‘B’ and “C”, the untranslated exon was present at −1366 to −1272 bp from the ATG initiation codon (Figs 3 and 4). This region shows high similarity with other teleost fish sequences as compared here through a phylogenetic tree presented in Supplementary Fig. S4 as well as values presented in Supplementary Table S2B. It is well established that the untranslated exon present at the 5’ regulatory region of a gene is transcribed into mRNA without translation [88]. However, several studies reported the importance of untranslated exon [89] towards the control of translation efficiency, stability of mRNA and also it influences gene expression efficiency of ORF [90,88]. In addition, Dvorak et al., (2019) [91] reported untraslated exon is having important role linked to animal evolution.

**Fig 4.**
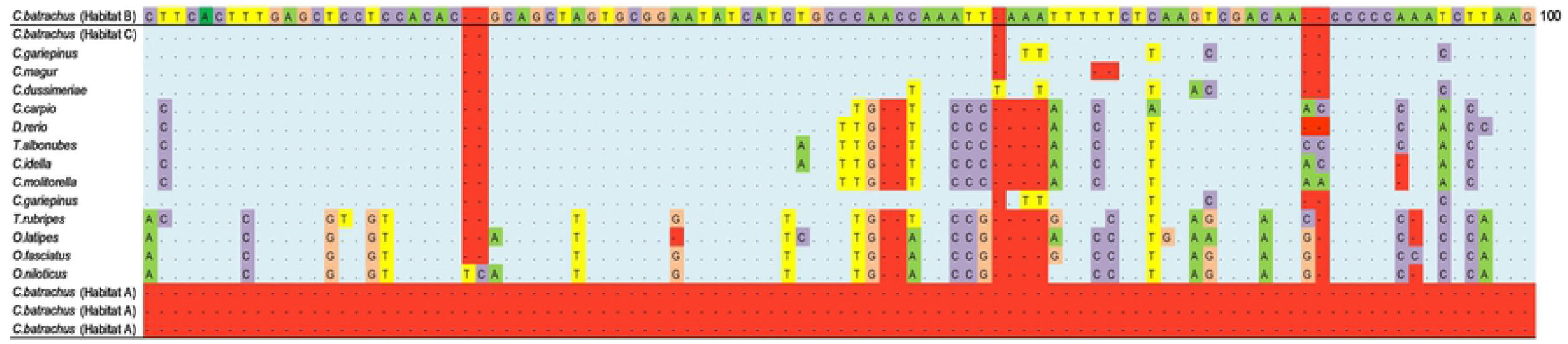
Sequence comparison of untranslated exon 1 from Catfish and representative teleostean β actin genes. Multiple sequence alignment of teleost β actin untranslated exon 1.Genbank accession no for each sequence is KJ722166.1 *(Clarias gariepinus),* M24113.1 *(Cyprinus carpio)*, EF026002.1 *(Danio rerio),* DQ241809.1 *(Cirrhinus molitorella),* M25013.1 *(Ctenopharyngodon idella),* EF026000.1 *(Tanichthys albonubes)*, U37499.1 *(Takifugu rubripes),* FJ975146.1 *(Oplegnathus fasciatus),* S74868.1 *(Oryzia slatipes),* AY116536.1 *(Oreochromis niloticus).* Dots indicates the identical nucleotide to *C. batrachus* from habitat “B” and “C”. Dashes (-) are gaps introduced for an optimal alignment.

ACpG island of 321 bp was predicted at position 1879-2199 within exon 3 (Fig 5). This sequence has 57 % G+C and 0.6 CpG ratio (observed/statistically expected, 0.6-0.8) in β actin gene using the database for CpG island following Kuo et al., [24]. The nucleotide composition was, A (21.95 %, n=277), T (32.81 %, n=414), C (22.90 %, n=289) and G (22.35 %, n=282) (S5 Fig). Interestingly, *C. batrachus* studied here does not have a CpG island at 5’ end of β actin gene near transcription start site (TSS). However, about 90% of housekeeping genes have promoter associated CpG islands near the transcription start site (TSS) [92–95]. Moreover, here the CpG island overlaps with exon3 which is referred to as an exonic region [96, 97] or an intragenic CpG island [98]. Consistent with findings of our study, Cuadrada et al., [99] noted that CpG islands are often species specific and unique in terms of their sequence and position related to the TSS. Also, our results suggest that the β actin promoter of *C. batrachus* is an “AT based promoter” with high A+T content (about 54 %) (S5 Fig) with CpG islands far from the TSS or in the 5’ regulatory region of gene [100]. This reduction of CpGs in the promoter or close to the TSS might have occurred due to nucleotide substitution and mutation in the process of evolution [101].

**Fig 5.**
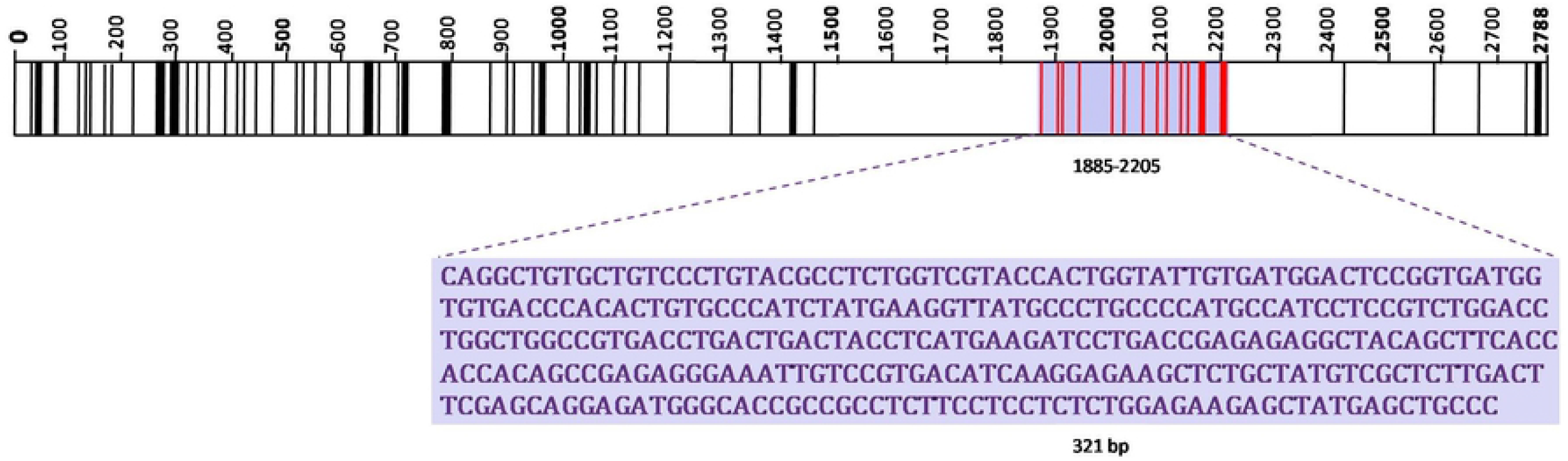
The schematic map of predicted CpG island in β actin genomic fragment of *C. batrachus*. Highlighted region shows the predicted CpG island. (Black colored vertical lines indicates the CpG sites. Red color line indicate the CpG sites in CpG island).

### Overview on organization of the β actin gene

Overall, the β actin gene described here display typical patterns of organization like other vertebrate cytoplasmic β actin genes. This gene has four completely translated exons (123 bp, 240 bp, 438 bp and 183 bp) and partial fifth translated exon (138 bp, here stop codon is not covered) with an open reading frame (ORF) of 374 amino acid residues (Fig 6 and S6 fig). The deduced amino acid sequence of the β actin protein described here has values for theoretical isoelectric point *(pI)* and molecular weight of 5.30 and 41 kDa, respectively.

**Fig. 6.**
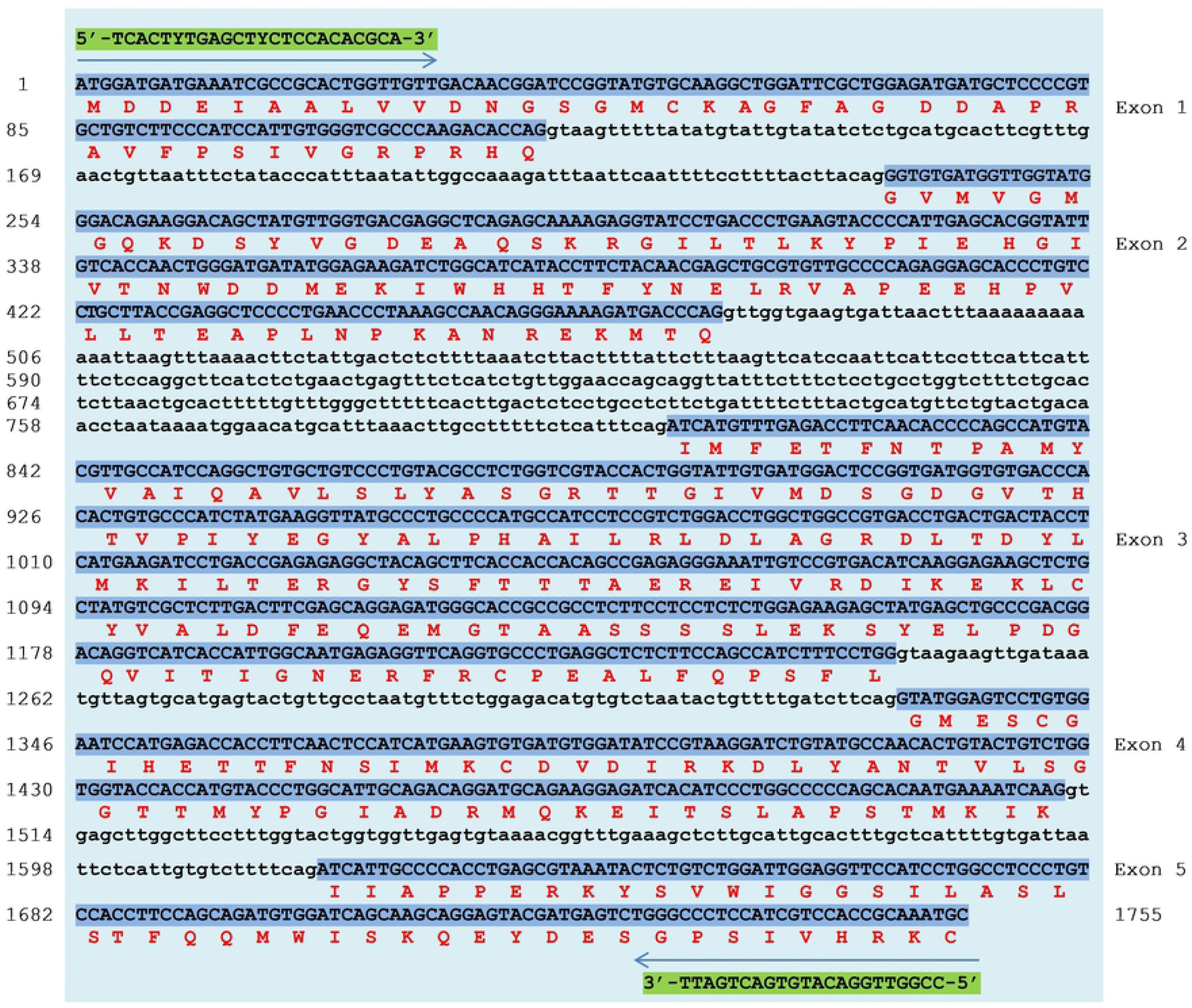
Complete nucleotide sequence and predicted amino acid residues of β actin gene. The capital and small letters indicate exons and introns respectively. The numbers on left side indicate the nucleotide position. The forward and reverse primer sequence used for amplification of β actin gene (arrow denotes the direction of primer).

The five translated exons were separated by four introns (111 bp, 330 bp, 83 bp and 106 bp), and the intron-exon boundaries were conserved following the GT/AG junction rule [102]. The genomic organization of β actin gene of *C. batrachus* was also defined by comparing it with the β actin gene from other teleost species. The translated exon sequences from *C. batrachus* showed 100% homology with earlier data from *C. batrachus* mRNA (GenBank Accession No. **LC379212.1**). Moreover, the *C. batrachus* β actin gene sequence shows 87.26%homology with *C. gariepinus* and 86.57% with *P. fulvidraco* (S2B Table). This implies that the genomic organization of *C. batrachus* is highly conserved among its populations as compare to other teleost β actin genes. To further deduce the relationship among *C. batrachus* individuals from populations as well as between four sister species, a phylogenetic tree was built using which supports this hypothesis (S7 Fig).

## Methods

### Ethical Statement

The Clarias species sampled for tissue biopsy are not protected under Indian wildlife protection act 1971. All fish are used as food fish and routinely caught and sold in local markets. No experimentations were conducted on live animals under this study.

### Sample collection

The tissue samples of *C. batrachus* were collected from three different habitats (S1 Table) with the help of local fisherman on a voluntary basis. From each habitat, approximately ten fish were sampled for tissue biopsy. Collected tissues were preserved in 1.5 mL airtight vials containing 85% ethanol. Samples were labeled and kept on ice in insulated boxes for transport to the laboratory for further processing.

### DNA extraction and quantification

Preserved tissue (10-15 mg) was used for nucleic acid purification by employing the Wizard genomic DNA kit (Promega, USA) and following the manufacturer’s protocol. Purified DNA was quantified (100 ng/μL), quality checked and stored at −20°C for further downstream applications.

### Amplification of the 5’ regulatory region of β-actin genes

The 5’ regulatory regions of the β-actin gene from all the Clarias species studied here were amplified with a set of degenerate primers derived from earlier studies [103–106]. Primers for PCR were synthesized by (IDT, India) on scale of 20Mm. The 25 μL PCR reaction was comprised of 1.0 μL (100 ng) genomic DNA, 2.5 μL 10X PCR buffer, 0.5 μL MgCl_2_, 3.0 μL (2.5mM) dNTPs, 1.0 μL (10mM) each primer, 0.4 μL of normal Kappa taq. (5U/μL) and17μL nuclease free water. The PCR cycling conditions were an initial denaturation step at 94°C for 2 min (one time) followed by 35 cycles of 50 sec at 94°C, 50 sec at 60°C, and 1 min at 68°C and a final extension at 68°C for 7 min. The amplified PCR products were visualized on 1% agarose gels for confirmation (Supplementary Fig. S1). Annealing temperatures used for each species were as follows: *C. batrachus* (60°C), *C. gariepinus* (57 °C), *C. magur* (55 °C) and *C. dussumieri* (56 °C).

### Amplification of the β-actin gene in different species

A novel pair of primers was also designed for amplification of the β actin gene based on conserved sequences of β-actin from *C. carpio*, *P. fulvidraco*, *C. gariepinus*, *O. latipe*, *C. idella* and *O. niloticus*. The PCR reaction (22 μL) comprises 1.0 μL of (100 ng) genomic DNA, 5.0 μL 5X GC PCR buffer, 0.5 μL, MgCl_2_, 4.0 μl dNTP (2.5mM), 1.0 μL (10mM) each primer, 1.0 μL taq. polymerase (2U/ μL) (ThermoFisherphusion high fidelity DNA Polymerase), 5.0 μL DMSO, and 4.5 μL of nuclease free water. The amplification was done with a touchdown PCR profile consisting of an initial denaturation step of 98°C for 30 sec, 35 cycles of 30 sec at 98°C, 50 sec at 58°C – 68°C and 2 min at 72°C and a final extension at 72°C for 10 min. The amplified PCR products were visualized on 1% agarose gels (S1 Fig).

The PCR products were purified to remove unincorporated primers and dNTP’s using the PCR purification kit (Purelink PCR purification kit, Invitrogen, USA). The purified products were processed for sequencing on a next generation sequencer platform @ 40X coverage (MiSeq, Illumina San Diego, California, USA).

### Sequence Analysis

#### Sequencing and preparation of read assembly

The obtained sequence reads of the β actin gene and its 5’ regulatory region were analyzed using the Galaxy Bioinformatics platform (http://www.bioinformatics.nl/galaxy) to check sequence read quality and assembly [107]. Sequence data is available through a link https://dataview.ncbi.nlm.nih.gov/object/PRJNA623463?reviewer=ia4fqq7jcgi94ngvimkcdchhre.

#### (b) BLASTn search and Annotation

The 5’ region and regulatory elements were predicted using GPMiner (http://gpminer.mbc.nctu.edu.tw/)[108–109]. The BLASTn search tool was also used to identify significant matches with 5’ regulatory regions of other β actin genes.

#### (c) Gene annotation

The FGENESH and GENSCAN gene finders were used for annotation. Predicted open reading frames (ORFs) and the inference of amino acid sequences was done using NCBI ORF finder tool (http://www.ncbi.nlm.nih.gov/gorf/gorf.html). Theoretical molecular weight value (kDa) and isoelectric point value (Ip) were calculated for the deduced amino acid sequences using the ExPAsy Prot Param tool (http://web.expasy.org/protparam/).

#### (d) Prediction of Cis-acting regulatory elements (CAREs) and CpG island

The transcription control elements TATA, CCAAT, GC and CArG boxes were predicted using the TFSEARCH tool (https://www.cbrc.jp/research/db/TFSEARCH.html). The *Cis*-acting elements in the 5’ regulatory region were identified using Cister and Cluster buster software (https://zlab.bu.edu/~mfrith/cister.shtml) [110]. The identification of CpG island was performed using the Data Base of CpG Islands and Analytical Tools (http://dbcat.cgm.ntu.edu.tw./) [111]. Nucleotide composition was calculated with MEGA program [112] and GC distribution along the promoter region was analyzed with the GC Content calculator (https://www.novoprolabs.com/) [113].

### Prediction of Untranslated exon

The prediction and phylogenetic analysis of untranslated exon was performed by comparing the β actin gene from other teleostean species. Publically available sequences were downloaded from NCBI (https://www.ncbi.nlm.nih.gov/). The sequences were aligned using Clustal-W and subjected to Maximum-Likelihood analysis with bootstrapping using the MEGA program [112]. For this analysis, sequences from mice *(Mus musculus)* and humans (*Homo sapiens)* were used as outgroups. The tree topologies were analyzed by bootstrap analysis with 1000 replicates.

## Conclusion

The 5’ regulatory region of the β actin gene of *C. batrachus* and other *Clarias* species has shown considerable nucleotide diversity and changes in promoter architecture with respect to addition or deletion of *Cis*-acting regulatory elements (CAREs). In some cases, our results also show that the organization of the β actin gene promoter does not follow a typical pattern in that this housekeeping genes shows reduced conservation in 5’ upstream sequence. We also show that some changes may be strongly linked to positive selection in dealing with pollution or other environmental challenges encountered by these organisms. These findings also support the idea that even housekeeping genes may play a critical role in sustaining organisms in diverse environmental conditions.

## Acknowledgement

Authors are thankful to DST INSPIRE, New Delhi, India for providing Senior Research fellowship to Ms. Deepali Sangale. We are also thankful to Director, CCMB, Hyderabad for extending their bioinformatics facility for data analysis. We sincerely thank all staff member and students at Paul Hebert Centre for DNA Barcoding and Biodiversity Studies, Aurangabad for their assistance in completing this work.

## Supporting information

**S1 Fig. Amplification of β actin gene and its 5’ regulatory region** (Size determination on 1% agarose gel)

**S2 Fig. Schematic map showing the genomic organization of 5’ regulatory region β actin gene from habitat “B” (1586bp)**

(Predicted transcription start site (TSS) is denoted with +1. First base of starting codon ATG was set as positon +1. The Exons (upper case letters in green color) and introns (lower case letters in black color) are shown in different boxes. *Cis*-elements including TATA Box, CAAT Box, CArG Box and GC Box are denoted in the figure panel. The highlighted part with aqua color indicate the diverse region of 556bp including untranslated exon1 (gray color) and binding site for additional 11 Cis-acting elements. Numbering for the nucleotide sequence position is given on left side

**S3 Fig. Schematic map showing the genomic organization of 5’ regulatory region β actin gene from habitat “C” (1572bp)** (Predicted transcription start site (TSS) is denoted with +1. First base of starting codon ATG was set as positon +1. The Exons (upper case letters in green color) and introns (lower case letters in black color) are shown in different boxes. *Cis*-elements including TATA Box, CAAT Box, CArG Box and GC Box are denoted in the figure panel. The highlighted part with aqua color indicates the diverse region of 541bp including untranslated exon1 (gray color) and binding site for additional 6 Cis-acting elements. Numbering for the nucleotide sequence position is given on left side.

**S4 Fig. Phylogenetic relationship of Clarias species and a representative teleostean fish based on untranslated exon 1.** Bootstrap values are shown at each node: dark circle > 90; dark square >50 and dark trangle <50.

**S5 Fig. Nucleotide composition of 5’ regulatory region of β actin gene.** [Full length-1262 bp, Average GC% (45%) and AT (54%) with distribution of nucleotide A(21.95 % n=277), T(32.81 % n=414), C(22.90% n=289), G(22.35 % n=-282)]

**S6 Fig. Comparison of amino acid sequences of clarias species β actin protein with other teleost.** The alignment was carried out using ClustulW. The highly conserved, identical residue indicated with “*” and less conserved residue was indicated with “:” symbols. The other protein were downloaded from NCBI *Pelteobagrus fulvidraco* (EU161065.2), *Cyprinus carpio* (M24113.1), *Ctenopharyngodon idella* (M25013.1, *Oryzia slatipes (S74868.1Oplegnathus fasciatus* (FJ975146.1), *Takifugu rubripes* (U37499.1 *Homo sapiens* (M10277.1) which was translated from its nucleotide sequences.

**S7 Fig. Neighbor-joining tree based on β actin gene sequence.** Bootstrap values (1000 replication steps) are shown at each node: dark circle > 95; dark square >90. Branch length is proportional to evolutionary distances. The scale bar indicate the evolutionary distance with respect to amino acid substitution per position in sequence.

**S1 Table.** Details of sampling station from major water bodies

**S2 Table.** NCBI BLASTn similarity search for β actin promoter and β actin gene

**S3 Table.** GC content of the promotor

